# Neuronal activity regulating the dauer entry decision in *Caenorhabditis elegans*

**DOI:** 10.1101/2024.09.30.615551

**Authors:** Sharan J. Prakash, Maedeh Seyedolmohadesin, Mark G. Zhang, Sarah M. Cohen, Shahla Gharib, Vivek Venkatachalam, Paul W. Sternberg

## Abstract

The model nematode *Caenorhabditis elegans* can choose between two alternative developmental trajectories. Larvae can either become reproductive adults or, under conditions of crowding or low food availability, enter a long-term, stress-resistant diapause known as the dauer stage. Previous studies showed that chemical signals from a secreted larval pheromone promote the dauer trajectory, and that their influence can be antagonised by increased availability of microbial food. The decision is known to be under neuronal control, involving both sensory and interneurons. To make an accurate decision, larvae must collect and compare complex patterns of environmental input over several hours of early larval development. The full composition of this circuit and the algorithm for decision-making are unknown. Here, we used cell-specific chemical silencing to systematically perturb several sensory and interneurons to further elucidate circuit composition. Our results suggest a role for gas-sensing neurons in regulating dauer entry. In addition, we quantitatively characterized the neuronal responses to food and pheromone inputs by measuring calcium traces from ASI and AIA neurons. We found that calcium in ASI increases linearly in response to food, and similarly decreases in response to pheromone, revealing a cellular site of antagonism between these key chemical inputs. Notably, the ASI response persists well beyond removal of the food stimulus, thus encoding a memory of recent food exposure. In contrast, AIA reports instantaneous food availability, and is unaffected by pheromone. We discuss how these findings may inform our understanding of this long-term decision-making process.

## Introduction

Phenotypic plasticity allows many organisms to undergo adaptive developmental and physiological changes in response to changing environmental conditions. The most dramatic forms of phenotypic plasticity are polyphenisms, in which individuals are able to choose between discrete alternative morphologies (Nijhout 2003). A well-known polyphenism in nematodes is the dauer diapause, an alternative developmental trajectory available to *Caenorhabditis elegans* larvae after the first (L1) larval stage (Fig. 1a) (Cassada and Russell 1975). Larvae enter dauer under adverse environmental conditions, and survive for several months without food (Klass and Hirsh 1976). When conditions improve, larvae can resume normal development at the L4 stage, without any effect on adult lifespan. The primary chemical inputs to the decision are microbial food signals, which promote adult development, and the dauer pheromone - a cocktail of secreted small molecules that promote dauer entry by providing a proportional signal of local crowding (Golden and Riddle 1982, 1984). Laser ablation and genetic studies established that entry into the dauer diapause is regulated by the nervous system (Bargmann and Horvitz 1991; Schackwitz et al. 1996; Zwaal et al. 1997). However, the full composition of the dauer circuit is unknown. To correctly anticipate future conditions and influence development accordingly, the nervous system must collect and compare fluctuating patterns of chemical inputs over the course of the early larval stages, emphasizing long-term trends. The complexity of this task suggests a distributed neural circuit capable of considerable plasticity and memory. Elucidating circuit composition is an important step in understanding the neuronal mechanisms of the decision-making algorithm. Recently, chemical silencing of individual neurons (Pokala et al. 2014) in combination with population assays were used to screen for neurons involved in dauer entry (Chai et al. 2022b). This study was the first to reveal a role for an interneuron (AIA). Gene knockout data also suggested that the activities of neuropeptide receptors expressed in a large number of neuron classes across the nervous system can modulate the decision (Chai et al. 2022a). In addition, microfluidics and calcium imaging experiments revealed that ASK and ADL sensory neurons are depolarized by pheromone (Chai et al. 2022b). Thus, a combined approach employing neuroanatomy, genetics, chemical silencing, behavior assays and calcium imaging can provide insight into neuronal regulation of dauer entry.

**Figure 1.**
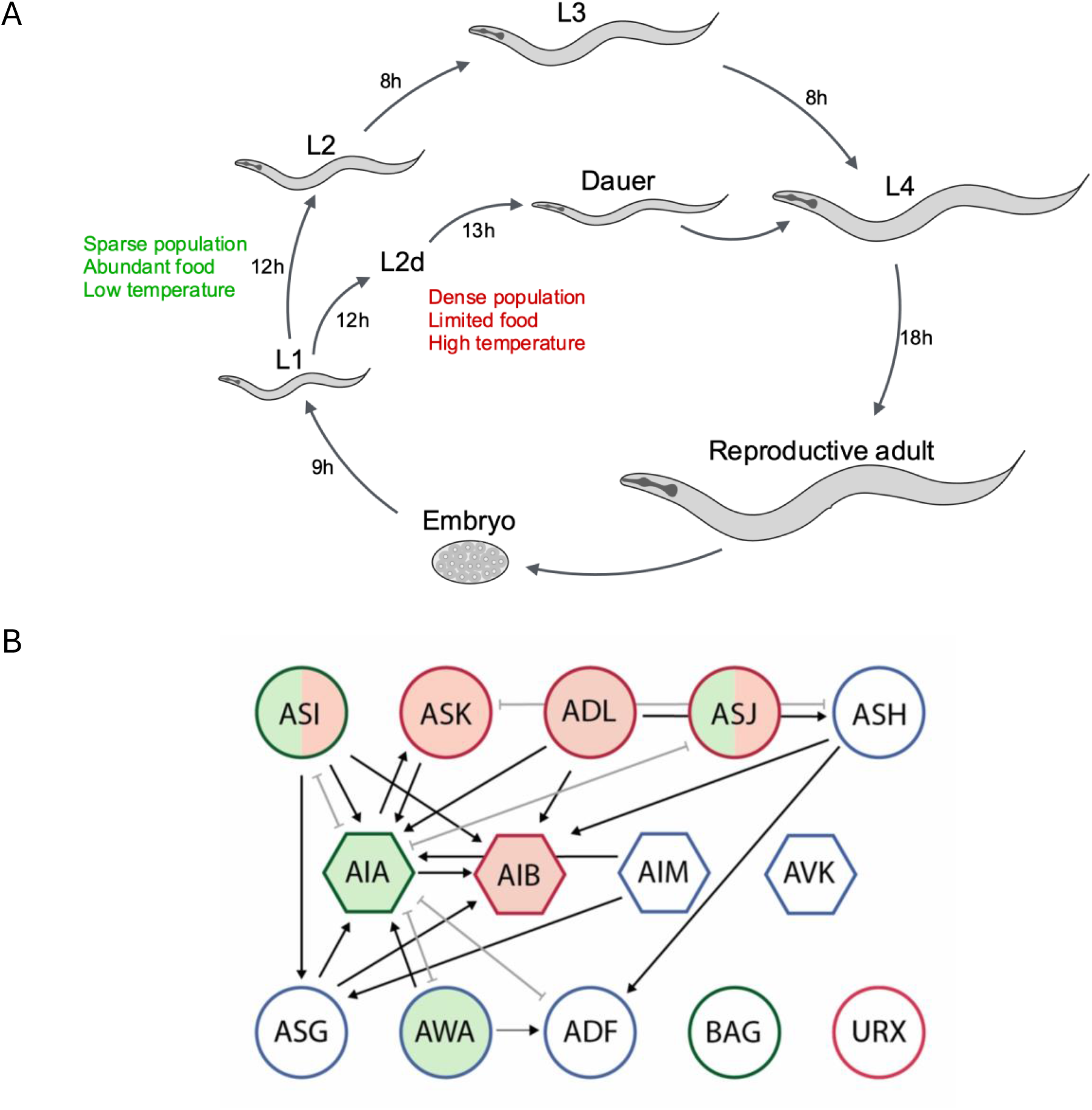
**A**. *C. elegans* life cycle. Larvae can enter the dauer stage after L1 under unfavourable conditions, and return to L4 when conditions improve. The progression to dauer involves an uncommitted L2d phase where larvae can escape dauer entry to L3, while committed dauers can only progress to L4, i.e. dauer is an alternative L3. **B**. Neural circuit diagram of all neurons mentioned in this paper. Sensory neurons are circles, and interneurons are hexagons. Outline colours refer to the valence of neuron depolarization on dauer entry (green = promotes adult development; red = promotes dauer entry; blue = no reported effect). Specifically, ASK and ADL promote dauer entry (Chai et al. 2022b), as do AIB (Chai et al. 2021), URX (this study) and ASJ (Schackwitz et al. 1996). Dauer-inhibiting neurons are ASI (Bargmann & Horvitz 1991; this study) and BAG (this study). Fill colours refer to response of the neuron to chemical inputs as evidenced by calcium imaging (green = depolarized by food; red = depolarized by pheromone; clear = no reported effect). Neurons depolarized by food are AWA (Larsch et al. 2015), AIA (this study), ASI (Chalasani et al. 2007; this study) and ASJ (Zhang et al. 2024). In the case of ASI and ASJ, the combined green and red fill indicates that food responses are inhibited by pheromone (this study and Zhang et al. (2024) respectively). Arrows refer to developmentally stable chemical synapses, while grey double-headed edges refer to gap junctions. Only connections including at least 3 chemical synapses and 2 gap junctions are included, for life stages embryo through L2. All circuit connectivity adapted from *nemanode*.*org* (Witvliet et al. 2021).

Here, we used chemical silencing and calcium imaging to identify additional neurons and study neuronal mechanisms involved in dauer entry. As expected, we found ASI inhibits dauer entry. We also found evidence for the BAG and URX sensory neurons in dauer regulation, while excluding several other sensory neurons and interneurons from the circuit (Fig. 1b). In addition, we compared the responses of ASI and AIA neurons to food and pheromone. We found that the calcium activity of ASI neuron displays a striking memory of food exposure, and is oppositely influenced by food and pheromone, while AIA reports the immediate availability of food, and is not responsive to pheromone. These results suggest specialisation of the sensory neuron ASI as a cellular site of antagonism between the key chemical inputs in the dauer decision. This contrasts with previous studies which describe the comparison of sensory inputs of opposite valence at the interneuron level (Ghosh et al. 2016; Gat et al. 2023).

## Results & Discussion

### Composition of the dauer circuit

The first studies to investigate the role of individual neurons in dauer entry used laser ablation, limiting their throughput (Bargmann and Horvitz 1991). To overcome this, we used chemical silencing via integrated transgenes in population assays. We found that silencing ASI promotes dauer entry (Fig. 2A), consistent with its known role as a food-sensor, and with patterns of expression of gene products that inhibit dauer formation (Murakami et al. 2001; Li et al. 2003; Cornils et al. 2011). Silencing ADF or ASG did not affect dauer entry (Fig. 2B, 2C) in contrast to previous laser ablation studies in which ASI, ASG and ADF were co-ablated (Bargmann and Horvitz 1991). ASH is receptive to a wide range of nociceptive stimuli, including hyperosmolarity and noxious chemicals (Hilliard et al. 2005), and inhibits ASI (Guo et al. 2015). We reasoned that the dauer decision may be receptive to output from ASH, but found no evidence for this (Fig. 2D).

**Figure 2.**
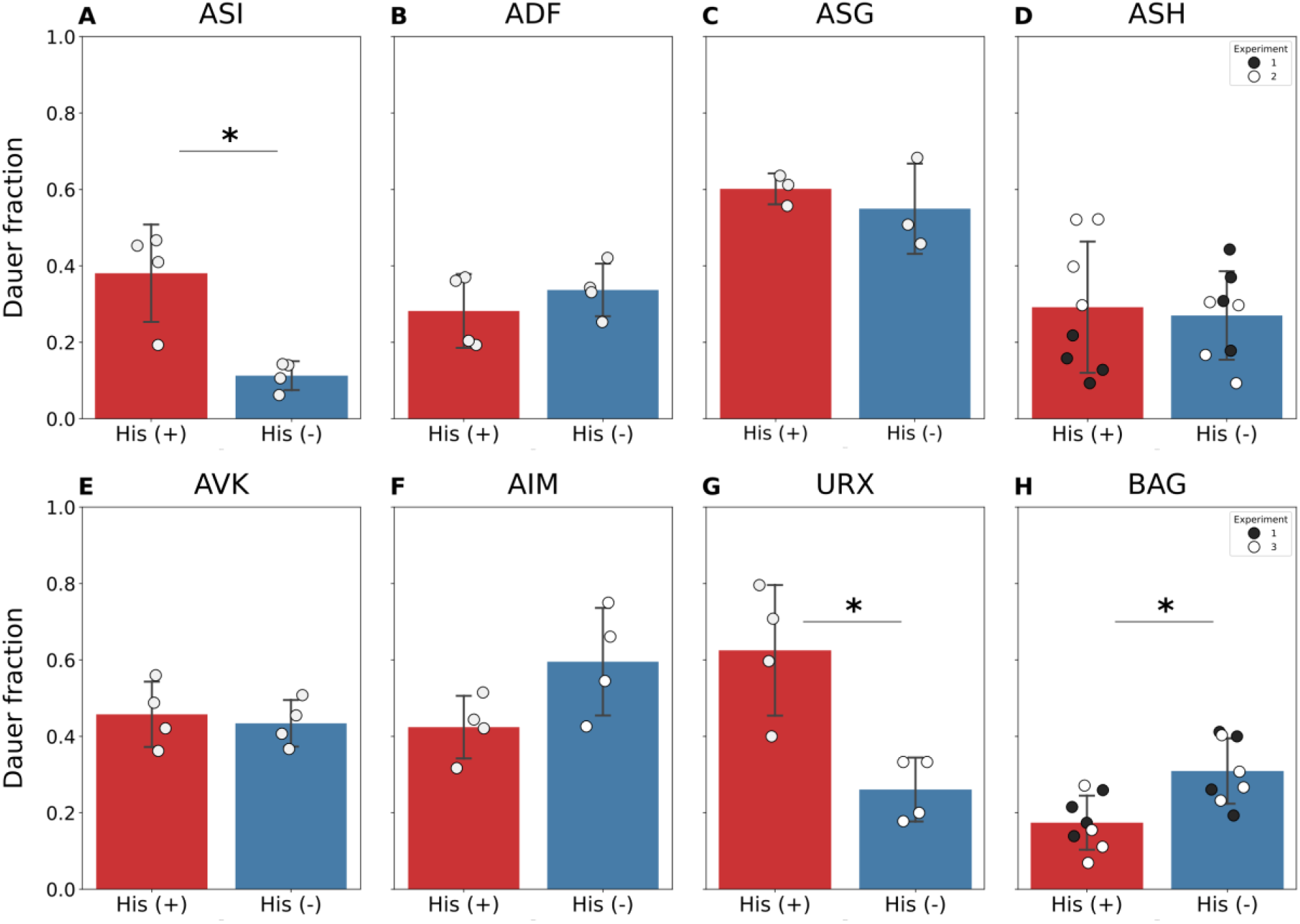
Screen for neurons involved in dauer entry. Animals expressing HisCl1 in individual neurons were grown on plates with (His (+)) and without histamine (His (-)). N=3-4 population assays (80-100 worms) per experiment. Stars (*) denote Student’s t-test (p<0.05). Data represented as mean with SEM.

We also tested BAG and URX, two sensory neurons receptive to gas stimuli. BAG mediates carbon dioxide avoidance (Hallem and Sternberg 2008) and is depolarized by pulses of carbon dioxide (Hallem et al. 2011; Bretscher et al. 2011; Carrillo et al. 2013) and downward shifts in oxygen concentration, while URX is depolarized by upward shifts (Zimmer et al. 2009). We found that inhibiting URX activity promotes dauer, while silencing BAG activity inhibits dauer (Fig. 2G, 2H). More experiments will be needed to clarify how these neurons are involved in dauer regulation. We note that these results are consistent with previous findings. A loss-of-function allele for the gene encoding the BAG-expressed neuropeptide FLP-17 showed decreased dauer entry (Lee et al. 2017). The cognate receptor for FLP-17, EGL-6, is expressed in URX, and both ablation of BAG and loss of FLP-17 increase the baseline fluorescence from URX-expressed calcium indicators (Hussey et al. 2018). URX was previously implicated in dauer by genetic evidence, in which loss-of-function alleles for URX-expressed neuropeptide receptors exhibited dauer entry phenotypes in each direction (Chai et al. 2022a). We also tested interneurons, including AVK and AIM (Fig. 2E, 2F), but found no evidence for an effect, despite a reported effect of AVK peptide output on transcription of dauer-inhibiting peptides in ASI (Une et al. 2022).

Overall, these results confirm the dauer-inhibiting role of ASI, and invite further investigation into the role of gas-sensing in dauer entry. At first glance, the absence of any effect from the interneurons tested contrasts with previous work, which suggested a large role for neuropeptide signalling activity across the brain in dauer entry (Chai et al. 2022a). In this work, loss-of-function alleles for 37 GPCRs, expressed widely in the brain, were found to either promote or inhibit dauer entry. However, for the majority of neuron classes, the set of expressed GPCRs included corresponding alleles with both dauer-promoting and dauer-inhibiting effects. Thus, the cell-specific GPCR expression patterns did not imply a consistent directional effect on the decision for most neuron classes, including AVK and AIM. We speculate that some interneurons may play a role in integrating cues from various sensory modalities or peptidergic indicators of internal state, each of which may modulate their effects on dauer entry over the course of the decision-making period.

### ASI and AIA respond differently to food and pheromone

An important question is to understand how the opposite effects of food and pheromone are reflected in antagonistic interactions at the molecular level, and the neurons in which this mechanism takes place. ASI expresses several pheromone receptors (Kim et al. 2009; McGrath et al. 2011; Park et al. 2012), is depolarized by soluble chemicals present in liquid OP50 solution (Chalasani et al. 2007), and synthesizes peptides that inhibit dauer formation in the presence of food, but not in the presence of pheromone (Ren et al. 1996; Murakami et al. 2001; Li et al. 2003). We hypothesized that this neuron may compare food and pheromone cues via changes in its electrical activity, with corresponding downstream effects on cell-autonomous transcription. The synaptic connectivity of interneuron AIA is also suggestive of a role in comparing food and pheromone signals. AIA is post-synaptic to the food sensor AWA (Larsch et al. 2015), as well as pheromone sensing ADL and ASK neurons, suggesting a neural pathway for its previously observed inhibition by pheromone. AIA inhibition promotes dauer entry (Chai et al. 2022b).

To investigate this, we delivered food and pheromone to worms immobilised in a microfluidic chip, and imaged the calcium dynamics of ASI and AIA neurons using GCaMP6s. We found that ASI responds strongly to OP50 supernatant, such that the amount of intracellular calcium increases with stimulus duration (Fig. 3A, left panel). It also displayed a striking memory of food exposure, remaining depolarized for at least 15 seconds after the stimulus is removed. Crude pheromone in isolation elicited no response in either neuron (Fig. 3A, middle panel). We found that adding crude pheromone to the food stimulus eliminated the food response in ASI, confirming our hypothesis (Fig. 3A, right panel). In contrast to ASI, AIA responded quickly to OP50 exposure (Fig. 3B, left panel) and showed no response to pheromone in isolation (Fig. 3B, middle panel). Previous work had suggested that pheromone inhibits spontaneous AIA activity (Chai et al. 2022b). Therefore we examined the possible inhibition of AIA by pheromone by mixing food and pheromone inputs. We saw no suppression of the AIA food response by pheromone (Fig. 3B, right panel), suggesting that AIA may not be highly sensitive to pheromone input from sensory neurons. Finally, by applying successive food and pheromone stimuli, we found that the depolarized state in ASI induced by food could be antagonised by subsequent exposure to pheromone (Fig. 3C).

**Figure 3.**
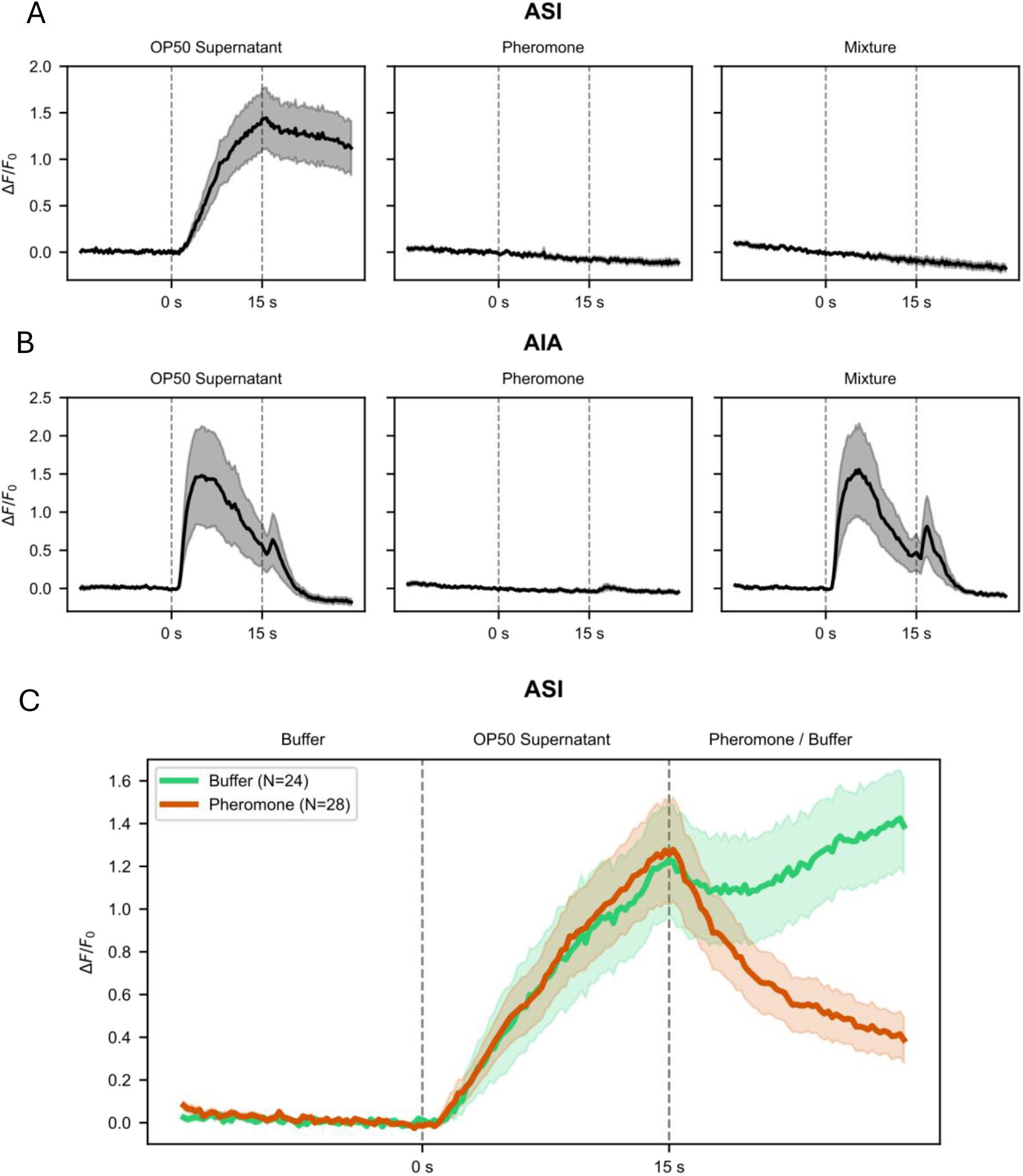
Calcium responses of ASI and AIA neurons to liquid inputs. L4 larvae were loaded into a microfluidic chip allowing exposure of sensory neurons each stimulus for 15 seconds. **A**. ASI exhibits slow dynamics in response to food (OP50 supernatant), with depolarization persisting beyond food removal (left panel), but is not depolarized by pheromone (middle panel). Mixture of food and pheromone eliminates the food response (right panel). **B**. AIA is rapidly depolarized by food (left panel), but not pheromone (middle panel). Mixture of crude pheromone and food results in a normal food reponse (right panel) **C**. ASI calcium response to bacterial food and pheromone applied in series. L4 larvae were exposed as before to OP50 supernatant for 15 seconds (both green and red lines), followed by exposure to crude pheromone extract for 15 seconds (red line only). Exposure to OP50 results in progressive depolarization which is reversed by addition of pheromone. In the control group, 15 seconds of OP50 exposure is followed by 15 seconds of buffer (green line). The depolarized state persists for at least 15 seconds after the removal of OP50.

The antagonistic effects of each input on ASI narrows the range of molecular mechanisms by which this regulation may take place to events upstream of neuron depolarization. For instance, receptors for food and pheromone components may have opposite effects on a common downstream ion channel, which in turn affects transcription of neuropeptide-encoding genes. In addition, ASI activity may regulate dauer entry via downstream neurons. Our finding that AIA’s food response is unaffected by pheromone suggests that its role in inhibiting dauer formation is primarily due to its role as a food-sensor, and suggests specialisation of ASI in the task of comparing food and pheromone signals. We speculate the primary role of AIA in dauer entry may be to transduce volatile signals produced by bacteria, sensed by pre-synaptic neurons such as AWA (Larsch et al. 2015). This also raises the question of how pheromone-sensing neurons ADL and ASK promote dauer entry. It will be informative to examine the effect of their activity on ASI and ASJ, for instance through combined optogenetics and calcium imaging experiments.

Our finding that ASI exhibits a memory of food exposure is especially exciting. A critical question in the study of the dauer circuit is to identify memory mechanisms that can bridge the short (seconds to minutes) timescales over which the relevant sensory inputs may fluctuate in duration and intensity, and the long timescale (hours) of the decision-making process. We hypothesize that the persistence of an ‘ON’ state in ASI represents such a bridging mechanism, allowing the production and secretion of peptides (such as DAF-28 and DAF-7) that promote adult development, and may themselves take part in persistent downstream signalling. Further, given that depolarization of ASI is antagonized by pheromone, the ASI response profile may facilitate an integration over time, i.e. the calcium response profile is proportional to the integral of historical differences between food and pheromone exposure. There is also a requirement for mechanisms that can bias the developmental trajectory in one direction or another, to ensure unambiguous commitment to one fate. The maintenance of an ‘ON’ state in ASI, promoting the continued production and secretion of dauer-inhibiting neuropeptides, could bias a developmental trajectory in the direction of reproduction based on recent history rather than instantaneous input alone, helping to ensure a binary outcome. In addition, the molecular or circuit mechanisms by which a long-lasting depolarized state can be maintained in a single neuron in the absence of continuous input deserves further investigation. One possibility is a positive feedback loop between two neurons. This motif appears elsewhere in dauer entry regulation, where positive feedback in the synthesis of Dafachronic Acid is vital to committing larvae to reproduction (Schaedel et al. 2012). We note that other mechanisms, such as changes in synaptic weight may also be important in bridging the timescales of neuronal activity and the physiological decision. Recent studies from our lab revealed similar calcium dynamics for the food response of ASJ neuron, as well as inhibition of the food response by pheromone (Zhang et al. 2024). The role of ASJ in dauer entry is less clear than for ASI. For instance, laser ablation experiments suggesting a dauer-promoting effect during the early larval stages (Schackwitz et al. 1996), and expression of dauer-promoting neuropeptide *ins-1* (Pierce et al. 2001). However, like ASI, ASJ is depolarized by food, and exhibits pheromone-regulated expression of dauer-inhibiting insulin *daf-28* (Li et al. 2003). The respective roles of ASI and ASJ in integrating food and pheromone cues in dauer entry and exit will require further study.

Going forward, studies of the dauer circuit will benefit from the use of calcium imaging from either whole brains, or neuronal subsets to describe the flow of sensory information through the nervous system. Along with long-term neuronal silencing experiments, these data can be tied to behavior. These data will help uncover the mechanisms and algorithms that regulate the decision to enter dauer.

## Methods and Materials

### Animal maintenance and strains

Animals were cultivated at 21°C on standard nematode growth media (NGM) plates seeded with *Escherichia coli* OP50 cultured in lysogeny broth (LB). All cGAL strains were generated by crossing UAS effector and GAL-4 driver strains generated by the Sternberg lab (Wang et al. 2017; Nava et al. 2023). The strain for imaging AIA (Chai et al. 2022b) and the strain used to image ASI (Lin et al. 2023) were previously published. The full list, origin, description and availability of strains used in this study can be found in Supplementary Table 2.

### Pheromone Preparation

Crude pheromone was prepared as described previously (Schroeder and Flatt 2014). Briefly, *C. elegans* were cultured in flasks with OP50 *E. coli* over several days until starved, with one additional round of feeding and starvation. Worms were separated from liquid by centrifugation and filtration. Crude pheromone was isolated by heating and ethanol extraction, and resuspended in water.

### Dauer assays

Dauer assays were performed as described previously (Lee et al. 2017; Chai et al. 2022b). On day 1, histamine chloride inhibition plates were prepared as follows. For each plate, 2ml of peptone-free Nematode Growth Medium (NGM) was mixed with either 6ul or 10ul of crude pheromone extract. For experimental plates, 30-40mg histamine dihydrochloride powder was first dissolved in 15ml NGM solution at 60°C, while control plates lacked histamine. Four plates were prepared per treatment for each experiment. One control plate lacking pheromone, but containing histamine, was also prepared for each experiment from the same histamine (+) NGM dilution. Roughly 100 L4 adults were picked onto new seeded plates, and a single colony of OP50 was used to inoculate LB at 37°C overnight. On day 2, OP50 was concentrated to 8% w/v in S-basal. 2ul of this OP50 was added to each plate to allow 80 adults to lay 70-90 eggs at 25°C. As a control, the line PS8720, expressing the His-Cl effector transgene under the *myo-3* promoter was transferred to the histamine (+) control plate. Successful inhibition results in cessation of locomotion in these animals. Remaining OP50 was heat killed at 95°-100°C for 10 minutes, and 18ul added per plate. Once dried, all plates were parafilmed and moved to 25°C for 72 hours. Dauer and non-dauer worms were counted for each plate.

### Statistical Analysis

All statistical analysis was performed with scipy. Jupyter Notebooks containing relevant code are available upon request. P-value were adjusted for multiple hypothesis testing via the Benjamini-Hochburg test (Benjamini et al. 2001), at a false discovery rate of 5% for are listed in Table 1. Original data are listed in Supplementary Table 1.

**Table 1.**
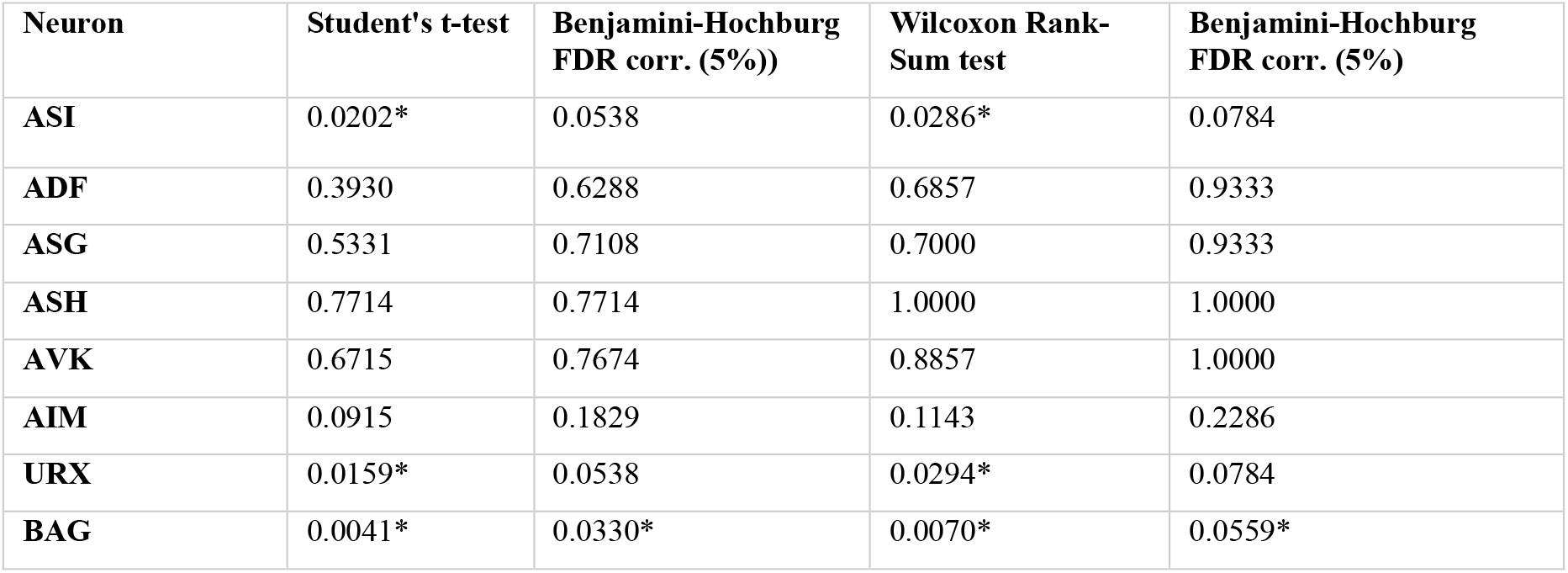
Statistical analysis of dauer decision-making assays after histamine-chloride inhibition. Stars denote p<0.05.

| Neuron | Student's t-test | Benjamini-Hochburg FDR corr. (5%) | Wilcoxon Rank-Sum test | Benjamini-Hochburg FDR corr. (5%) |
| --- | --- | --- | --- | --- |
| ASI | 0.0202* | 0.0538 | 0.0286* | 0.0784 |
| ADF | 0.3930 | 0.6288 | 0.6857 | 0.9333 |
| ASG | 0.5331 | 0.7108 | 0.7000 | 0.9333 |
| ASH | 0.7714 | 0.7714 | 1.0000 | 1.0000 |
| AVK | 0.6715 | 0.7674 | 0.8857 | 1.0000 |
| AIM | 0.0915 | 0.1829 | 0.1143 | 0.2286 |
| URX | 0.0159* | 0.0538 | 0.0294* | 0.0784 |
| BAG | 0.0041* | 0.0330* | 0.0070* | 0.0559* |

### Microfluidic device fabrication

The Venkatachalam lab designed a 2-layer microfluidic chip capable of delivering sequences of stimuli with a worm trap suitable for housing worms at L4 larval stage. The chip was designed in AutoCAD software, and sent to Artnet Pro Inc. for photomask printing. Photolithography in a clean room was performed on a silicon wafer to make the 2-layer mold from the photomask. For the first layer, which included the worm trap, SU-8 2025 was spin coated on the silicon wafer at 4000 rpm to achieve 25 μm thickness. For the second layer, the same photoresist was spin coated at 1250 rpm for a thickness of 70 μm. Polydimethylsiloxane (PDMS) was poured over the mold and cured on a 90^°^C hotplate to solidify. Each PDMS chip was then punched with a 1 mm biopsy punch and was bonded to a cover slip using a handheld corona treater.

### Calcium imaging

L4 stage animals were assayed and placed in the microfluidic device. For imaging of AIA, a strain expressing GCamp6s specifically in AIA was used (PS9111). For imaging of ASI, a strain expressing GCamp6s in all sensory neurons was used (ZM10104). For each experiment, *E. coli* OP50 was cultured overnight in LB, and its supernatant was collected. Crude pheromone extract was diluted to a concentration of 2.5% (v/v) in either buffer (H2O) or OP50 supernatant. Each stimulus was delivered to the animal’s nose for a duration of 15 seconds, followed by either buffer (H_2_O) or pheromone. Fluorescence was recorded with a spinning disc confocal microscope (Dragonfly 200, Andor) and a sCMOS camera (Photometrics Kinetix). The fluorescence was captured from GCaMP6s at a rate of 10 ms per 1.0 μm z-slice, with 25 z-slices per volume and 4 volumes per second. To extract calcium activity from the recorded data, we performed the following steps: 1. Background intensity was subtracted from each recorded volume. 2. The center of the ROI was annotated at one timepoint, and then the center was tracked throughout the entire recording using the Zephir tracking algorithm (Yu, 2022). 3. For ASI neurons, the ROI was defined as the neuronal nucleolus, and for AIA neurons, the ROI was defined as the processes located in the gap junction. 4. Average pixel intensity from each ROI was calculated. ΔF/F0 was computed, where F0 was defined as the average intensity during the 5-second window preceding stimulus delivery.

## Declarations

### Data & Materials Availability

All data generated or analyzed during this study are included in this pre-print and its supplementary information files. A complete list of strains and corresponding genotypes is provided in the supplementary materials.

## Competing Interests

The authors declare that they have no competing interests.

## Author Contributions

S.J.P. and M.S. are co-first authors. P.W.S, V.V., and S.J.P. conceived and designed the study. S.J.P. conducted crude pheromone extraction, and dauer assays. M.S. conducted calcium imaging experiments. S.J.P. performed the data analysis for dauer assays. M.G.Z. and S.M.C. provided crude pheromone reagents for imaging experiments. S.J.P. and S.G. generated new cGAL strains. S.J.P. wrote the manuscript with editorial assistance from M.S., V.V. and P.W.S. M.S. and S.J.P. prepared all the figures. All authors approved the manuscript.

## Funding

S.J.P. was supported by NIH Grant U01 NS111697, M.G.Z. was supported by a NIH Grant F31 NS120501-01, P.W.S. was supported by a Bren Professorship and by a NIH Grant R24-OD023041. V.V. and M.S. were supported by a Burroughs Wellcome Fund Career Award at the Scientific Interface and NIH R01 NS126334.

## Acknowledgements

We thank all members of the Sternberg lab at Caltech for their feedback during the course of the project.

## Notes

### Competing Interest Statement

The authors have declared no competing interest.

### Summary of Updates

Updated funding information, author attributions, and abstract.

